# LSTrAP-*denovo*: Automated Generation of Transcriptome Atlases for Eukaryotic Species Without Genomes

**DOI:** 10.1101/2023.03.05.530358

**Authors:** Peng Ken Lim, Marek Mutwil

**Affiliations:** School of Biological Sciences, Nanyang Technological University, 60 Nanyang Drive, Singapore, 637551, Singapore

**Author notes:** Corresponding author: Marek Mutwil, School of Biological Sciences, Nanyang Technological University,60 Nanyang Drive, 637551, Singapore, Singapore.

## Abstract

**Motivation:** Despite the abundance of species with transcriptomic data, a significant number of the species still lack genomes, making it difficult to study gene function and expression in these organisms. While *de novo* transcriptome assembly can be used to assemble protein-coding transcripts from RNA-sequencing (RNA-seq) data, the datasets used often only feature samples of arbitrarily-selected or similar experimental conditions which might fail to capture condition-specific transcripts.

**Results:** We developed the Large-Scale Transcriptome Assembly Pipeline for *de novo* assembled transcripts (LSTrAP-*denovo*) to automatically generate transcriptome atlases of eukaryotic species. Specifically, given an NCBI TaxID, LSTrAP-*denovo* can (1) filter undesirable RNA-seq accessions based on read data, (2) select RNA-seq accessions via unsupervised machine learning to construct a sample-balanced dataset for download, (3) assemble transcripts via over-assembly, (4) functionally annotate coding sequences (CDS) from assembled transcripts and (5) generate transcriptome atlases in the form of expression matrices for downstream transcriptomic analyses.

**Availability and Implementation:** LSTrAP-*denovo* is easy to implement, written in python, and is freely available at https://github.com/pengkenlim/LSTrAP-denovo/.

**Supplementary Information:** Supplementary data are available in the forms of supplementary figures, supplementary tables, and supplementary methods.

## 1. Introduction

In recent years, the number of publicly available RNA-sequencing (RNA-seq) data uploaded to the International Nucleotide Sequence Database Collaboration (INSDC; Burgin et al., 2023; Sayers et al., 2022; Tanizawa et al., 2023) databases has accumulated rapidly (Katz et al., 2022; Lowe et al., 2017). This is due to the advancements in next-generation sequencing (NGS) technology that can sequence short-reads (reads) with increasing affordability and higher throughput (Levy & Myers, 2016). This has not only added to the existing transcriptomic data of model species, allowing for more powerful secondary analyses (Julca et al., 2023; Lim, Zheng, et al., 2022) but has importantly, also provided first-ever transcriptomic data for many species without a genome. The expression data could be used to quantify gene expression and to study gene function and transcriptional regulation (Julca et al., 2023; McDermaid et al., 2019; Usadel et al., 2009). To exemplify, there are 1677 *Viridiplantae* species (as of Feb 2023; Table S1) with at least 10 RNA-seq experiments on the European Nucleotide Archive (ENA; Burgin et al., 2023). However, only 528 (31.5%) out of the 1677 species have a representative genome assigned by the National Center for Biotechnology Information (NCBI) Genome database (https://www.ncbi.nlm.nih.gov/genome/; NCBI Resource Coordinators, 2016). This disparity is largely attributed to the relative ease with which transcriptome data can be generated via RNA-seq, compared to the difficulty in assembling and annotating high-quality complete genomes, especially for organisms with large, repetitive, complex, and/or allopolyploid genomes (Giani et al., 2020; Kyriakidou et al., 2018; Zhang et al., 2022).

Fortunately, various *de novo* transcriptome assemblers have been developed to assemble transcripts directly from RNA-seq reads without genomic references (Bushmanova et al., 2019; Grabherr et al., 2011; Liu et al., 2016; Peng et al., 2013; Schulz et al., 2012; Xie et al., 2014) and can be used as pseudoalignment references to expression quantification. Importantly, transcripts assembled from this method are not affected by genome quality. As a result, *de novo* transcriptome assembly methods are useful to assemble protein-coding transcripts of species without reference genomes or with poor-quality genomes.

The current *de novo* transcriptome assembly approaches often use in-house RNA-seq datasets capturing similar experimental conditions, developmental phase, and organ/tissue type (for multicellular organisms), which can fail to capture condition-specific (for e.g. tissue-specific) transcripts (Alsamadany, 2020; Han et al., 2022; Joudaki et al., 2023; Prochetto et al., 2023). While the transcriptome coverage (experimental conditions/sample type) of an RNA-seq dataset can be improved by using public RNA-seq data (Gilbert, 2019), the types of samples to include in the RNA-seq dataset are often not well-defined. Moreover, due to the differences in the quality of publicly available data, the *de novo* transcriptome assembly workflows need to comprise multiple steps, such as quality control of RNA-seq data (Bolger et al., 2014; Chen et al., 2018), adaptor trimming of reads (Bolger et al., 2014; Chen et al., 2018), transcriptome assembly and transcript annotation (Cantalapiedra et al., 2021; Finn et al., 2011; Nawrocki & Eddy, 2013; Teufel et al., 2022). These steps involve multiple tools which require considerable bioinformatics expertise to implement, making transcriptome assembly inaccessible to non-bioinformaticians.

To address this paucity, we developed the Large-Scale Transcriptome Analysis Pipeline for *de novo* assembled transcripts (LSTrAP-*denovo*). Given an NCBI taxonomic ID (NCBI TaxID), the pipeline automatically downloads publicly available RNA-seq experiments (accessions) from ENA, assembles transcripts, and generates transcriptome atlases for eukaryotic species.

Specifically, LSTrAP-*denovo* can (1) automatically filter out low-quality or wrongly curated RNA-seq accessions, (2) select RNA-seq accessions via unsupervised machine learning to construct a sample-balanced dataset for *de novo* transcriptome assembly, (3) assemble transcripts via over-assembly, (4) functionally annotate coding sequences (CDS) from assembled transcripts and (5) generate transcriptome atlases in the form of expression matrices. This is achieved via 5 connected scripts (termed modules; Figure 1) to be sequentially executed via the CLI (command line interface) in Linux-based operating systems or Windows Subsystem for Linux (WSL).

**Figure 1.**
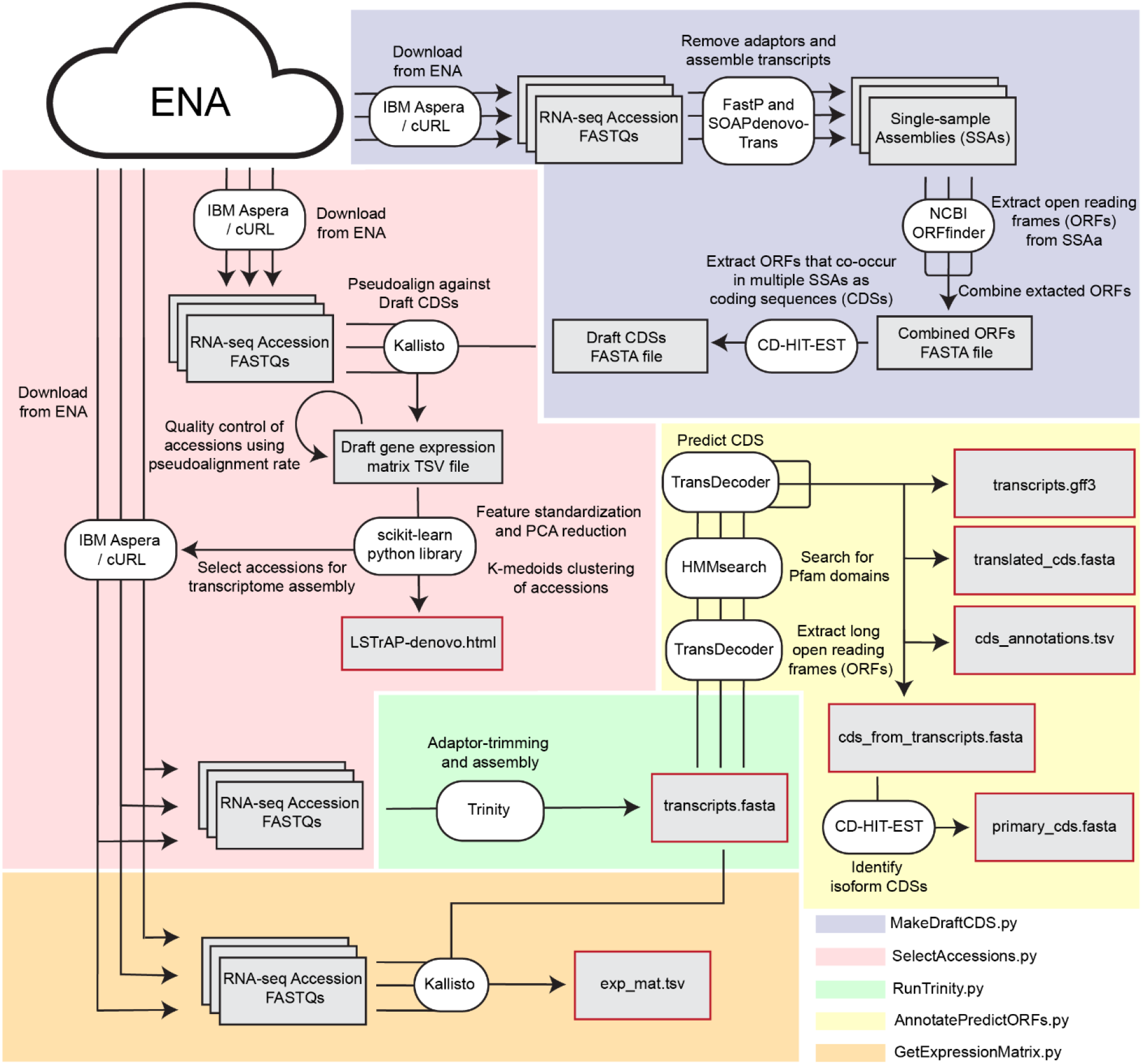
Simplified schematic of LSTrAP-*denovo*. The pipeline consists of 5 modules to be executed sequentially by individual scripts (MakeDraftCDS.py, SelectAccessions.py, RunTrinity.py, AnnotatePredictORFs.py, and GetExpressionMatrix.py demarcated by blue, red, green, yellow, and orange respectively). Application Programming Interface (API) access to European Nucleotide Archive (ENA) is represented by a cloud icon. Arrows indicate the flow of information (input and output) through the command-line interface (CLI) tools/functions from the SciKit-learn python library (represented by white text bubbles). Parallelized processes are represented by parallelized arrows while intermediate and output files generated by the pipeline are represented by gray text boxes outlined in black and red borders, respectively.

## 2. Implementation and Algorithm

### 2.1 Generation of Draft CDS via over-assembly

The analysis starts with the MakeDraftCDS.py module (Figure 1, blue demarcation) of LSTrAP-*denovo*. For a given species specified by NCBI TaxID, LSTrAP-*denovo* will first download a randomized subset (10 accessions by default) of RNA-seq accessions from ENA, remove read-adaptors using FastP (Chen et al., 2018) and then assemble every accession individually using SOAPdenovo-Trans (Xie et al., 2014). SOAPdenovo-Trans was chosen due to its relatively low memory usage and fast runtime (Hölzer & Marz, 2019; Zhao et al., 2011), allowing for the parallelization of assembling transcripts. In order to maintain similar read coverage for every RNA-seq accession, accessions are capped to the same size (1.5 GB gunzipped by default) prior to adaptor removal and assembly.

Next, a collection of open reading frames (ORFs) are generated for every accession (henceforth referred to as Single-sample Assemblies [SSAs]) by using ORFfinder (https://www.ncbi.nlm.nih.gov/orffinder/). The SSAs are then unified into a single over-assembly (Figure 2A), where CD-HIT-EST (W. Li & Godzik, 2006) clusters and identifies similar ORFs that share 98% global sequence similarity.

**Figure 2.**
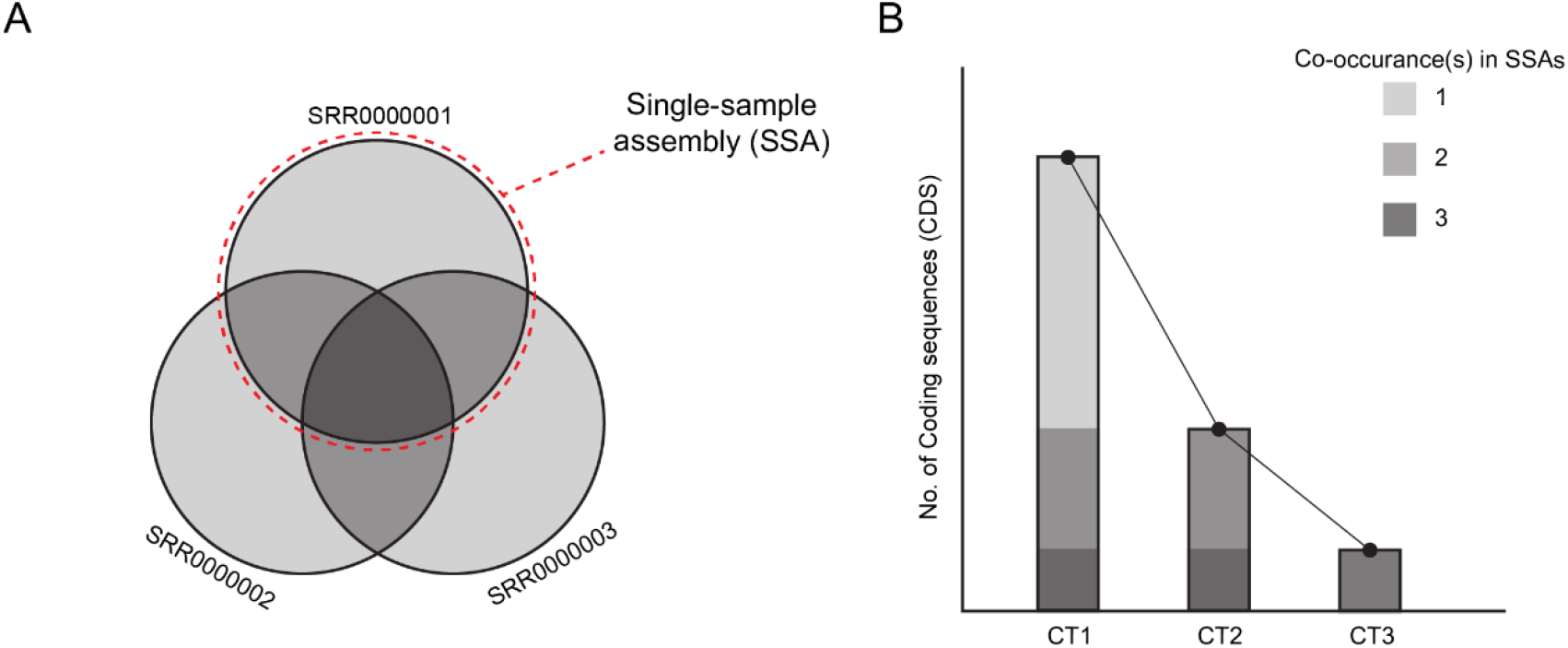
Method to obtain highly-probable Draft Coding Sequences. (A) Over-assembly is made by unifying ORFs from three single-sample assemblies (SSAs). Each circle of the Venn diagram represents an SSA assembled from one RNA-seq accession (denoted by mock SRA accession identifiers). Groups of ORFs that co-occur in multiple SSAs are represented by overlapping regions. (B) A consensus approach to extract Draft CDS from over-assembly. The combined line and the stacked-bar plot show how the use of different CTs can yield a different number of Draft CDSs

Finally, the ORF over-assembly is refined further into highly-probable Draft coding sequences (CDS), by extracting only ORFs that co-occur in multiple SSAs (Figure 2B). Specifically, an ORF is considered as “co-occurring” if highly similar ORFs (98% global sequence similarity; determined by CD-HIT-EST) are found in multiple SSAs. ORFs with a co-occurrence above a set threshold (consensus threshold [CT]) is considered Draft CDSs (Figure 2B). The expected number of Draft CDS is set as the median number of ORFs across all SSAs. MakeDraftCDS.py automatically detects a CT that yields the number of Draft CDSs that is the closest to the expected number, but the user has the option to specify a custom CT.

### 2.2 Pseudoalignment of RNA-seq accessions against Draft CDS for quality control and expression estimation

This part of the pipeline, along with 2.3 and 2.4, is executed by the SelectAccessions.py module (Figure 1, red demarcation). LSTrAP-*denovo* will download all RNA-seq accessions for the species of interest, and pseudoalign each RNA-seq accession against the Draft CDSs generated in section 2.1 using the Kallisto (Bray et al., 2016, p.). The distribution of pseudoalignment rates (PR) from RNA-seq accessions (in the form of the % pseudoaligned metric reported by Kallisto) is used to identify and exclude undesirable RNA-seq accessions for transcriptome assembly (e.g., non-coding RNA-seq or RNA-seq data of a different organism); RNA-seq accessions are deemed undesirable if PRs are below the lower bound of the normal distribution (Q1 - (1.5 × IQR)) and/or if PR is <20%. The estimated Transcripts per million [TPM] expression values are used to construct a Draft expression matrix.

### 2.3 Unsupervised clustering of RNA-seq accessions

First, LSTrAP-*denovo* standardizing expression values of each RNA-seq accession within the Draft expression matrix to fit a standard normal distribution (zero-mean and unit variance). The normalized matrix is transformed using Principal Component Analysis (PCA) (Yeung & Ruzzo, 2001) into a feature matrix that describes RNA-seq accessions in the first 100 principal components. If the number of RNA-seq accessions that passed QC in 2.2 is below 100, the number of principal components is set to be equal to the number of RNA-seq accessions.

Next, the feature matrix is used to embed RNA-seq accessions in a latent space (henceforth referred to as Draft expression space [DES] for brevity) for K-medoids clustering (Mohammed & Abdulazeez, 2017; Van der Laan et al., 2003) based on Euclidean distance. Since the K-medoids clustering algorithm requires specifying the number of expected clusters (*k*) *a priori*, LSTrAP-*denovo* constructs multiple K-medoids clustering iterations through a range of *ks* and selects the best *k* based on silhouette coefficient (Rousseeuw, 1987). To be conservative in estimating the optimal k, LSTrAP-*denovo* selects the lowest k that falls within 0.01 of the highest recorded silhouette coefficient amongst all clustering iterations.

### 2.4 Selection and download of RNA-seq accessions to establish a sample-balanced dataset

LSTrAP-*denovo* will first sort RNA-seq accessions in each cluster based on their distance from the cluster medoid. Next, LSTrAP-*denovo* will sequentially select and download paired-end RNA-seq accessions for each cluster until a specified read depth is reached for the cluster (starting from accessions with DES embeddings closest to the cluster medoid) to form a cluster-specific library. As with 2.1, the read depth of each cluster-specific library is approximated using file sizes of RNA-seq accessions (gunzipped FASTQ files).

### 2.5 Assembly and annotation of transcripts

The assembly of transcripts is executed by the RunTrinity.py module (Figure 1, green demarcation). By default, LSTrAP-*denovo* will adopt an over-assembly approach for *de novo* transcriptome assembly where Trinity (v2.14; Grabherr et al., 2011) is used to assemble each cluster-specific library downloaded in 2.3. Assembled transcripts from each cluster are combined into one FASTA file (transcripts.fasta; Figure 1) and CD-HIT-EST (W. Li & Godzik, 2006) is used to remove redundant transcripts (global similarity ≥ 98%). Alternatively, users can also use LSTrAP-*denovo* to feed all downloaded RNA-seq data from 2.3 into Trinity to generate a single transcriptome assembly.

The annotation of transcripts is executed by the AnnotatePredictORFs.py module (Figure 1, yellow demarcation). LSTrAP-*denovo* first uses TransDecoder (https://github.com/TransDecoder/TransDecoder) to extract translated open reading frames (ORFs) within transcripts. Next, Hmmsearch (Finn et al., 2011) scans the extracted ORFs against known Pfam (Blum et al., 2021) hmm profiles. TransDecoder then predicts CDSs (outputted as cds_from_transcripts.fasta; Figure 1) in assembled transcripts by evaluating the feasibility of CDSs based on a coding score as well as the presence of Pfam domains within the CDS (https://github.com/TransDecoder/TransDecoder). LSTrAP-*denovo* also provides an annotation file (cds_annotations.tsv; Figure 1) that maps identifiers of predicted CDSs to detected Pfam domains and their associated Interpro accessions (Blum et al., 2021) and Gene-ontology (GO) terms (Ashburner et al., 2000) to facilitate downstream analyses.

D*e novo* transcriptome assemblies are known to yield many transcript isoforms from the same genomic locus (Gilbert, 2019) that can be challenging to use as transcriptomic references to reliably estimate gene expression using pseudoalignment. To remedy this, LSTrAP-*denovo* generates a subset of primary CDSs (cds_from_primary_transcripts.fasta; Figure 1) from the total pool of predicted CDSs. LSTrAP-*denovo* achieves this by first identifying the groups of transcript isoforms by clustering predicted CDSs based on global sequence similarity ≥ 98% using CD-HIT-EST (W. Li & Godzik, 2006). This allows the selection of a primary CDS with the best coding score (as calculated by TransDecoder) for every isoform group.

### 2.6 Generation of transcriptome atlas

This step of the pipeline is executed by the GetExpressionMatrix.py module (Figure 1, green demarcation) of LSTrAP-*denovo*. In a fashion similar to 2.2, LSTrAP-*denovo* will download all RNA-seq accessions for the species of interest and pseudoalign them to CDSs of primary transcripts (cds_from_primary_transcripts.fasta) generated in 2.5 using Kallisto (Bray et al., 2016). The estimated relative expression values (in the form of Transcripts per million [TPM]) of RNA-seq accessions from pseudoalignment are used to construct a final gene expression matrix for downstream transcriptomic analyses (exp_mat.tsv; Figure 1).

## 4. Results

### 4.1 LSTrAP-*denovo* generates highly probable Draft CDSs using a novel consensus strategy fueled by over-assembly

Here, we present an automated and data-driven approach in CDS and expression matrix construction via LSTrAP-*denovo* (see 2.1 to 2.4) where a small subset of public RNA-seq data of a species of interest is initially used to assemble Draft CDSs as a pseudoalignment references for RNA-seq accession reads. This facilitates two aspects of dataset construction. Firstly, accessions undesirable for transcriptome assembly (e.g. ncRNA-seq accessions, misclassified RNA-seq accessions from other species) can be flagged and disregarded based on mapping rate as they are expected to contain reads that align more poorly to CDSs as compared to the majority of the other RNA-seq accessions (Proost et al., 2017). Secondly, the expression levels of Draft CDSs can be used as features to cluster RNA-seq accessions according to their transcriptome profiles (see 2.3 for feature preprocessing and clustering methodologies) to identify the most diverse set of RNA-seq accessions, that captures organ-, tissue- and cell-specific transcripts (see 2.4 for how accessions are selected).

To make this process automatic, the pipeline first randomly selects a number of RNA-seq accessions to assemble Draft CDSs. However, this random selection may include undesirable RNA-seq accessions that can negatively affect the accuracy of assembled CDSs. Over-assembly methods involve combining transcripts generated from multiple assemblers, as misassembled transcripts are unlikely to be found in more than one assembly (Gilbert, 2019; Voshall et al., 2021; Yu et al., 2020). We developed a similar over-assembly strategy specifically to exclude misassembled, exogenous, and fragmented transcripts that are assembled from read data of bad quality. Instead of combining assemblies generated from different assemblers, ORFs of randomly selected RNA-seq accessions are independently assembled to form SSAs (see 2.1) and subsequently combined. Using a consensus approach, only ORFs that co-occur in a set number of SSAs (consensus threshold [CT]; automatically determined) are designated as Draft CDSs. Since erroneously-assembled and exogenous ORFs are unlikely to appear more than once in SSAs, the Draft CDS likely represent bona fide coding sequences.

To evaluate the performance of this consensus approach, we used *LSTrAP-denovo* to generate Draft CDSs from 10 randomly selected accessions in four species (see supplementary methods S1). We compared these ORFs to ones that were assembled from one combined library of the 10 selected accessions (Combined library ORFs). Contig recall scores and Nucleotide precision scores calculated using REF-EVAL (B. Li et al., 2014) were used to assess the veracity of ORFs with reference CDSs as ground truth (see supplementary methods S2). For a given set of extracted ORFs, Contig recall is calculated as the number of reference CDSs recovered at full-length (99%) in the set of ORFs divided by the total number of reference CDSs while Nucleotide precision can be calculated as the sum of nucleotide lengths of ORFs within the set with perfect alignment to reference CDSs divided by sum of nucleotide length of all ORFs (B. Li et al., 2014). Because Nucleotide precision measures precision at the nucleotide level, non-canonical ORFs extracted from fragmented transcripts nested within true CDSs were accounted for in true positives, thus limiting false positives to ORFs originating from misassembled, exogenous, or non-coding transcripts.

The optimal consensus threshold (CT) used to extract co-occurring ORFs in the consensus strategy to generate Draft CDSs were automatically determined as described in 2.1. Across all four species, Draft CDSs generated using the consensus strategy achieved a higher Contig recall (1.54 to 2.50 fold change) and Nucleotide precision (1.11 to 4.53 fold change), when compared to Combined library ORFs, despite comprising fewer sequences (Figure 3). The disparity of Nucleotide precision scores between Draft CDSs and Combined library ORFs was the largest for *C. albicans* where the former has a precision of 82.4% while the latter only has a precision of 18.2%. Although precision seemingly improves the least for *A. thaliana* (i.e. 83.9% in Combined library ORFs to 92.9% in Draft CDSs; Figure 3), the improvement is still substantial. When considering the amount of false sequences reduced at the nucleotide level, the Combined library ORFs of *A. thaliana* contains 5.20 million nt of false sequences, but this number is reduced by 2.84 fold in its Draft CDSs (1.84 million nt; data not shown). The reduction in false sequence data is even more pronounced in *C. albicans* (34-fold reduction; 35.44 million nt vs 1.04 million nt; data not shown) as Combined library ORFs outnumber Draft CDSs more than 10 fold (71598 vs 7084; Figure 3).

**Figure 3.**
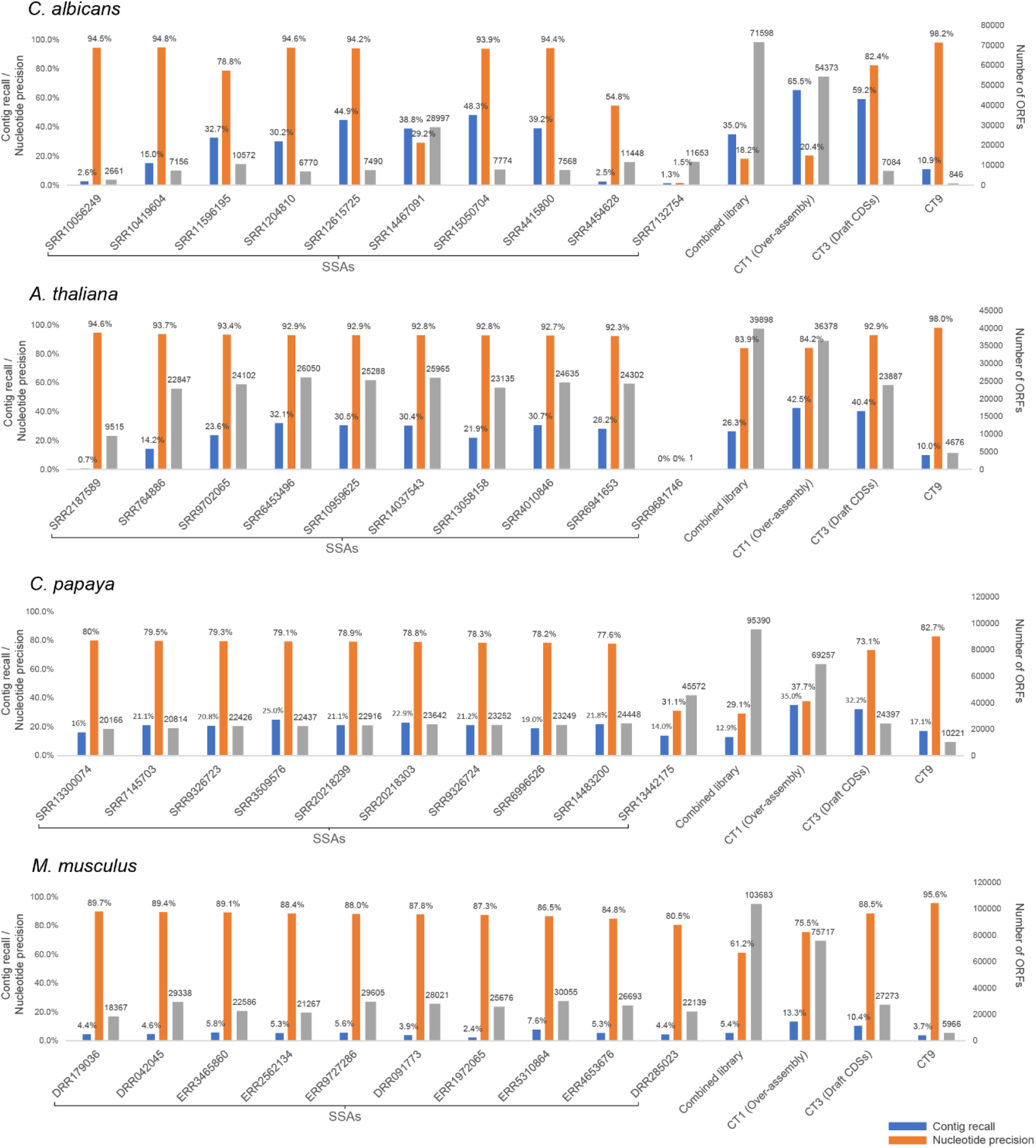
Evaluating precision and recall of Draft CDSs generated by LSTrAP-*denovo*. Bar charts show the precision (orange) and recall (blue) of ORFs from SSAs and Over-assemblies generated by LSTrAP-denovo. Precisions and recalls are represented by Nucleotide precision and Contig recall scores generated by the REF-EVAL module of DETONATE when aligned against designated reference CDSs. Gray bars indicate the number of ORFs and correspond to the secondary (right) axis.

To better understand how the consensus approach is effective in improving the accuracy of Draft CDSs, we investigated the precision and recall scores for ORFs extracted from independently assembled SSAs (SSA ORFs), and ORFs pooled at different CT thresholds. Scores calculated for *C. albicans* SSA ORFs display the greatest variability (Figure 3). In particular, precision scores for SSA ORFs of accessions SRR14467091, SRR4454628, and SRR7132754 were much lower than other *C. albicans* SSA ORFs (Figure 3), with SRR7132754 SSA ORFs carrying 98.5% of unmatched nucleotide sequence data (Nucleotide precision of 1.5%; Figure 3) within its 11,653 ORFs. Through analyzing the metadata of these accessions (Table S2) and their linked publications (Kämmer et al., 2020; Ost et al., 2021; Sieber et al., 2018), we found that these accessions hold RNA-seq data for the tissue samples of mice/human infected with *C. albicans*, which might result in the presence of host RNA sequences in the SSA ORFs of these accessions, and therefore, have low precision scores. Similarly, We have also identified an outlier SSA with low precision (SRR13442175; 31.1%; Figure 3) in *C. papaya* that likely contains exogenous fungal transcripts as the accession comprises RNA-seq data of plant sample infected with *Colletotrichum brevisporum* (https://www.ncbi.nlm.nih.gov/sra/?term=SRR13442175; Table S2). It is likely that the low precision scores of the Combined library ORFs and Over-assembly ORFs in C. papaya and C. albicans were caused by the presence of these accessions. Consequently, the consensus approach improves precision drastically because exogenous sequences are not likely to occur in multiple independently-sampled SSAs. Additionally, in *A. thaliana*, SRR2187589, and SRR9681746 SSAs have scores that were different from other SSAs. SRR2187589, a TAIL-seq accession (table S2), resulted in an SSA with a very low recall (0.7%; Figure 3). However, despite the low recall of SRR2187589, its precision is still relatively high (94.6%) while miRNA-seq accession SRR2187589 yielded only one ORF (Figure 3). As such, their negative impact on the precision of the Over-assembly ORFs and Draft CDSs is minimal.

Furthermore, assembling accessions individually and pooling ORFs recovers more true CDSs than assemblies generated from combined libraries. The recalls observed in Over-assembly ORFs and Draft CDSs are higher than recalls of Combined library ORFs across all species (Figure 3) despite the sequencing depths of the Combined libraries being one order of magnitude higher than the sequencing data from individual accessions (see 2.1). For example, for *C. albicans* we observed Contig recall of 35% for combined, 65.5% for Over-assembly ORFs, and 59.2% for assembled Draft CDSs. The exact reason for this remains to be elucidated but we believe that there exist certain biases within different libraries (resulting from differences in read length, read representation, expression pattern, etc.) that might favor the assembly of certain transcripts and these biases can disappear when libraries were all combined for assembly.

Overall, we have shown that the consensus approach employed by LSTrAP-*denovo* is able to generate highly probable Draft CDSs (with high precision and improved recall) from a set of randomly selected accessions of varying quality.

### 4.2 Pseudoalignment against Draft CDS reveals undesirable RNA-seq accessions

The publicly available RNA-seq data consists of samples of varying quality. Furthermore, some samples can be mislabeled and might actually represent data that are not suitable for transcriptome assembly. Over the years, the repertoire of available NGS-based transcriptome sequencing has rapidly expanded beyond just RNA-seq (of total RNA / mRNA) to include many other RNA-seq variations such as miRNA-seq, scRNA-seq, and Ribo-seq. However, INDSC databases do not have individual sections dedicated to these variations. Instead, sequencing data from all these RNA-seq variations might be classified under the umbrella of transcriptomic data. Because RNA-seq data available on INDSC databases are user-uploaded without strict requirements on the format of accompanying metadata, there exist many unannotated accessions of these RNA-seq variations on INDSC databases. Accessions of these RNA-seq variations are not useful for *de novo* transcriptome assembly and it is important for the pipeline to identify these undesirable RNA-seq accessions. Previous approaches removed undesirable accessions based on how well their reads map to their transcriptome references during pseudoalignment (Lim, Davey, et al., 2022; Tan & Mutwil, 2020; Villanueva et al., 2022). Although high-quality transcriptome references are not available for species without genomes, we believe that pseudoalignment of accessions against Draft CDSs can similarly be used to identify undesirable accessions. In this study, this is supported by the high correlation (r> 0.89) between pseudoalignment rates reported from pseudoalignment of accessions against Draft CDSs and the references CDSs (Figure S2).

To test the effectiveness of this approach, we downloaded random RNA-seq accessions for each species and pseudoaligned them against their respective Draft CDSs using LSTrAP-*denovo*. Approximately 1000 accessions were processed for model species (984 for *C. albicans*, 991 for *A. thaliana*, and 985 for *M. musculus*, and all available RNA-seq accessions for *C. papaya* (309). Next, we annotated the accessions by analyzing the metadata (Table S3 S4, S5, and S6), and classified them into 4 different categories (DNA-seq, ncRNA-seq, RNA-seq variants, and Exogenous contamination; Table S7). We observed that across all species, pseudoalignment rates of accessions follow a bimodal distribution (Figure 4; blue bars) where a small but significant portion of accessions have low pseudoalignment rates. Not surprisingly, the majority of ‘ncRNA-seq’ accessions (Figure 4, orange bars) belong to these poorly mapping accessions. ‘RNA-seq variants’ accessions (only found for *A. thaliana* and *M. musculus)* are largely distributed within the rate range of 0% −10% in *A. thaliana* but are evenly dispersed in *M. musculus*. These accessions are predominantly accessions holding scRNA-seq data (Table S4 and S5), and contain reads with unique molecular identifiers (UMI) and cell barcodes that can interfere with their pseudoalignment against transcriptomic references, hence explaining the low rates of these accessions observed in *A. thaliana*. In *C. albicans*, a known human pathogen, we found many RNA-seq accessions that might originate from human /mouse tissue (Exogenous contamination; Figure 4, red bars; Table S3), which also showed poor mapping.

**Figure 4.**
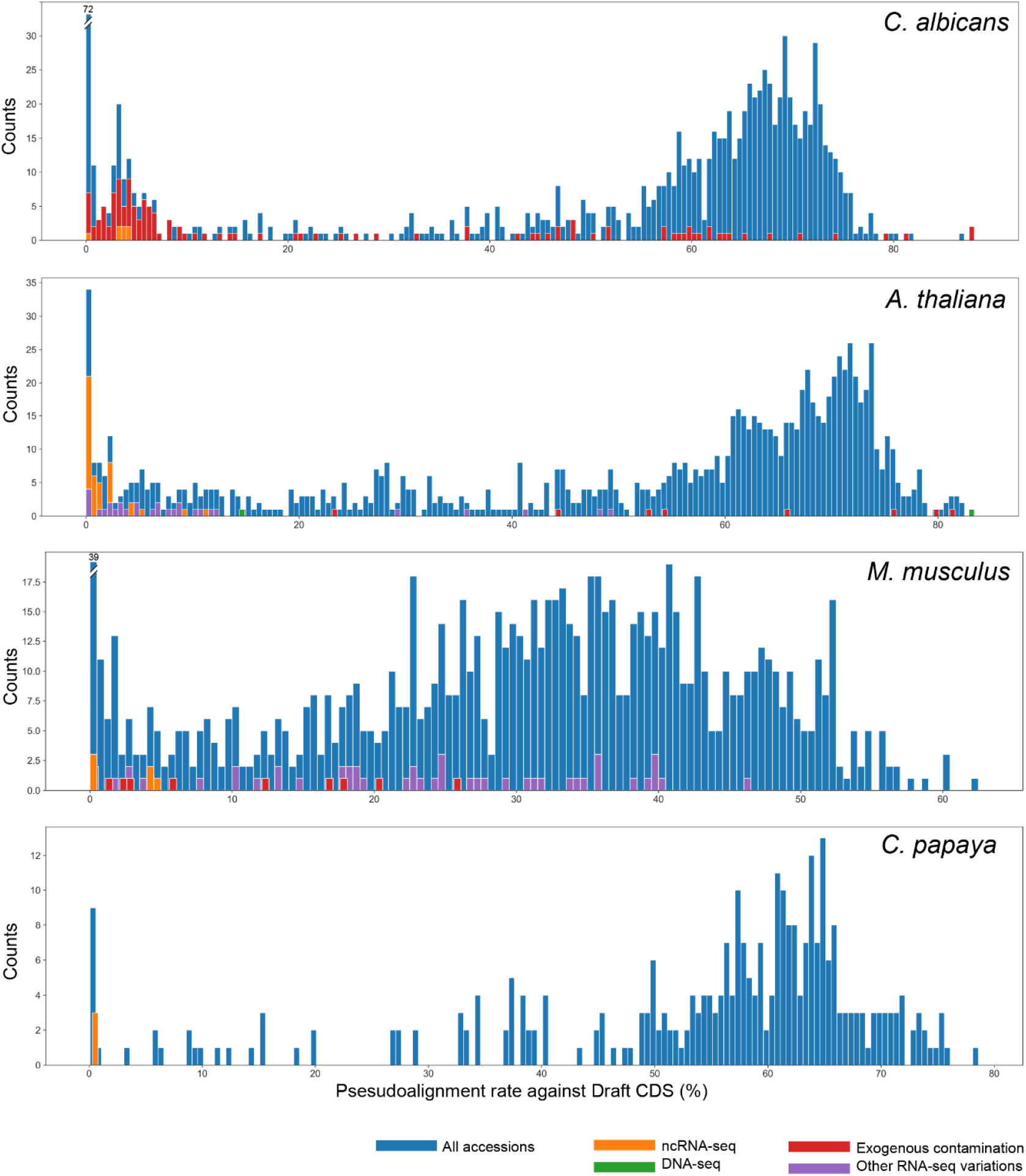
Pseudoalignment rates of different types of sequencing data. The histogram shows the distribution of Kallisto pseudoalignment rates (blue bars) of accessions from *C. albicans* (984), *A. thaliana* (991), C. papaya (309), and *M. musculus* (985) pseudoaligned to their respective Draft CDSs. The pseudoalignment rates of accessions capturing ncRNA-seq, DNA-seq, and cases where biological samples might contain exogenous contamination (inferred from title metadata) are represented as purple, orange, green, and red bars.

Overall, pseudoalignment rates can be used to flag undesirable accessions capturing incorrect types of sequencing data (e.g., miRNA-seq or rRNA-seq) or contamination by other organisms. While this approach might not be able to identify scRNA-seq data reliably (as seen for *M. musculus)*, their impact on *de novo* transcriptome assembly might be minimal given that k-mers from randomly generated UMI and cell barcodes have much lower counts than k-mers from transcripts. For all species, RNA-seq accessions with a pseudoalignment rate of < 20% were excluded in the downstream dataset selection process.

### 4.3 Sample selection with unsupervised clustering

The inclusion of diverse sets of RNA-seq data for the transcriptome assembly is likely to increase the detection rate of organ-, tissue-, cell- and condition-specific genes that might be missed if using a limited diversity of data. LSTrAP-denovo uses unsupervised clustering to identify samples that capture diverse transcriptomic profiles.

Since our assembled Draft CDS only used a small subset of available data, we observed relatively low Contig recall scores, and CDSs from exogenous, misassembled, and fragmented transcripts, which resulted in low Nucleotide precision scores (Figure 3, Table S8). For instance, in Draft CDSs of *M. musculus*, only 9,669 CDSs (out of 27,273; Contig precision of 35.5%; Table S8) accounting for 9,695 reference CDSs (out of 93,224; Contig recall of 10.4%; Table S8) are assembled to full length. Furthermore, 16,580 Draft CDSs do not align perfectly to any reference CDSs with a region coverage of >= 90% (Table S8), and likely coming from exogenous, misassembled, and fragmented transcripts.

To identify the diverse set of RNA-seq accessions that can be used to assemble a comprehensive set of coding sequences, we used k-medoids clustering (Van der Laan et al., 2003), where the euclidean distance between RNA-seq accessions represents a measure of similarity in global gene expression (Figure 2).

To evaluate how well the draft CDS can identify the clusters of samples, we compared clusters obtained from pseudoaligning the RNA-seq data to the draft CDS to clusters obtained from pseudoaligning to reference CDS (see supplementary methods S4 and S5). LSTrAP-*denovo* selects RNA-seq accessions closest to cluster medoids to establish a sample-balanced dataset (25% of accessions of each cluster closest to medoid; see supplementary methods S4) with a roughly equal amount of reads from each cluster (defined by the total file size of accession libraries). For a fair comparison of clusters obtained from the draft and reference CDS, we limit embedding comparisons to RNA-seq accession selected from the draft CDS. The similarity of draft and CDS reference clusters was evaluated with the V-measure metric (Villanueva et al., 2022), and the significance of the similarity was obtained with a Monte Carlo-based hypothesis testing (see supplementary methods S6).

For every species, we observed statistically significant similarities (P= 0.00001; Figure 5) between draft (DES) and reference (RES) CDS clusterings with observed similarity (in descending order) of 0.855 in *C. albicans*, 0.812 in *C. papaya*, 0.766 in *A. thaliana* and 0.705 in *M. musculus* (Figure 5). Importantly, the draft and reference clusterings were able to identify nearly identical clusters (Figure 5; asterisk bars) despite using different independently determined *k* (expected clusters). We observed 4, 5, 2, and 6 identical clusters between the draft and reference CDS for *C. albicans, A. thaliana*, *M. musculus*, and *C. papaya*, respectively (Figure 5, identical clusters indicated by *). Since the clusters obtained by draft and reference CDS are highly similar, we conclude that the Draft CDSs generated from a small number of samples can be used to select diverse RNA-seq accessions for the final transcriptome assembly.

**Figure 5.**
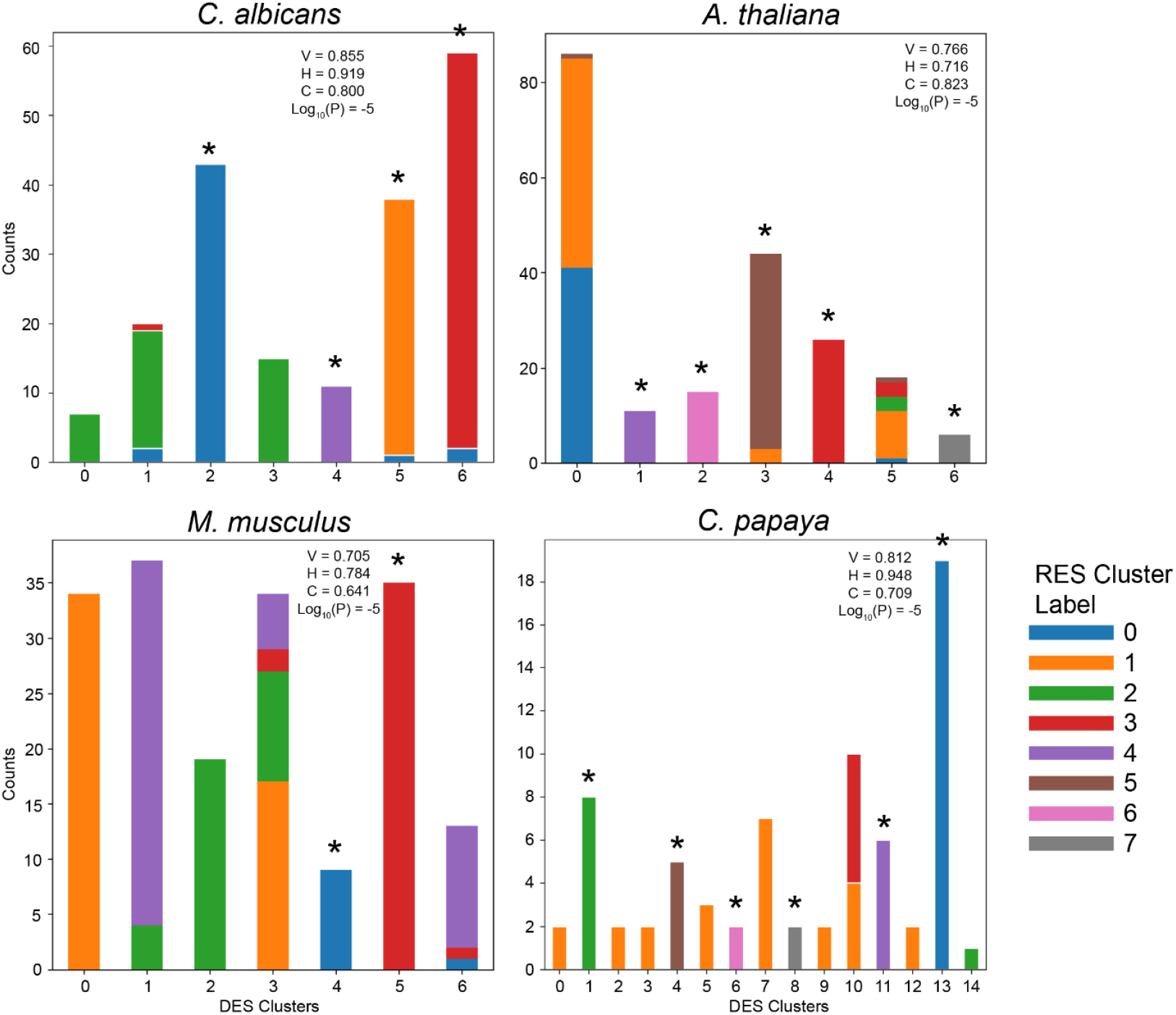
K-medoids cluster assignments of RNA-seq data when pseudoaligned to the draft and reference CDSs. Stacked-bar plots compare the similarity between draft and reference cluster assignments. The x-axis bars correspond to Draft CDS clusters while colors correspond to clusters obtained when mapping to the reference CDS. The optimal number of clusters (k) for k-medoids clustering was selected based on the highest silhouette score within 5 ≥ k ≤ 20. Optimal k for clustering for Draft clusters are 7, 7, 7, and 15 while optimal k for clustering of reference clusters are 5, 8, 5, and 8 for *C. albicans, A. thaliana, M. musculus, and C. papaya*, respectively. V, H, and C correspond to V-measure, homogeneity, and completeness scores, respectively, where the reference CDS clusters are treated as ground truth. Asterisks indicate near identical clusters observed for the draft and reference CDS.

### 4.4 Co-expression analysis of *C. papaya* transcriptome atlas reveals co-regulation of MEP and carotenoid biosynthesis pathway

To demonstrate how LSTrAP-denovo can be used to retrieve biologically-relevant information, we compared the assembled transcriptome and the genome of *C. papaya* (Papaya1.0). The genome available on NCBI was partially assembled in 2008 (Ming et al., 2008) covering only a portion of the complete genome. Consequently, the transcriptome sequence information for C. papaya is incomplete, as reflected in the analysis of Benchmarking universally conserved orthologs (BUSCOs) provided by NCBI, with only 80.8% of BUSCO completeness score (completeness score; Figure 6A; https://www.ncbi.nlm.nih.gov/data-hub/genome/GCF_000150535.2/).

**Figure 6.**
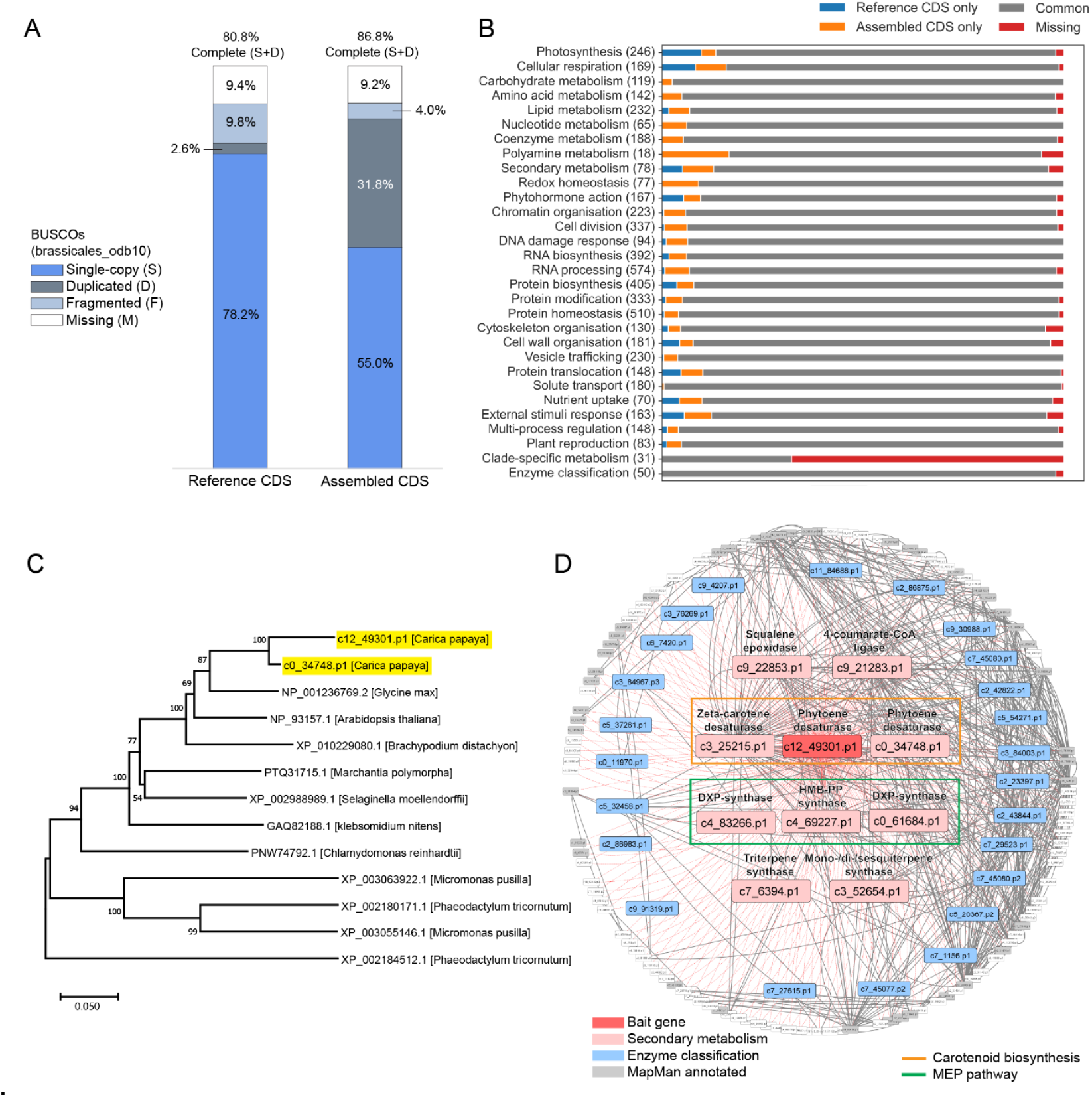
Transcriptome assembly and co-expression analysis of *C. papaya*. (A) BUSCOs analysis of assembled and reference *C. papaya* CDSs. Stacked bars percentages of single-copy, duplicated, fragmented, and missing *Brassicales* BUSCOs (odb10). (B) Comparison of functional MapMan bins in *C. papaya* assembled and reference CDSs. MapMan bins are categorized into functional groups according to their first-level descriptions (first-level bins). The barplot shows the proportion of MapMan bins within each group that are shared in the reference and assembly (gray), unique to reference (blue), unique to assembly(orange), and missing (not detected in both reference and assembled CDSs; red) (C) Phylogenetic analysis of assembled phytoene desaturases of *C. papaya* (highlighted blue and yellow) and phytoene desaturases of other *Viridiplantae* species. The phylogenetic tree was constructed by the neighbor-joining method using the MEGA11 program. Bootstrap values (1000 resamplings) are displayed next to branches. Protein accession identifiers of phytoene desaturases and their corresponding species (in square brackets) are displayed in the nodes. The tree is drawn to scale, with branch lengths in the same units as those of the evolutionary distances used to infer the phylogenetic tree. The evolutionary distances were computed using the p-distance method and are in the units of the number of amino acid differences per site. All positions containing gaps and missing data were eliminated (complete deletion option). (D) The local co-expression neighborhood of “bait gene”, *c12_49301.p1*. Network visualization features co-expression relationships between *c12_49301.p1* and the top 150 primary transcript CDSs most co-expressed with *c12_49301.p1*. Thin red edges connect *c12_49301.p1* to its 150 most co-expression CDSs while thick edges connect CDSs that are co-expressed with a PCC ≥ 0.85. With the exception of *c12_49301.p1* which is colored deep red, CDS nodes are colored based on their assigned first-level MapMan bins.

The first two modules of LSTrAP-*denovo* (MakeDraftCDS.py and SelectAccessions.py; Figure 1) discovered 15 clusters of RNA-seq accessions (Figure 5) and automatically downloaded a sample-balanced dataset (see 2.3 and 2.4) consisting of pair-end libraries of 34 accessions (Table S9). Through the use of a *de novo* transcriptome assembly method based on over-assembly (see 2.5), LSTrAP-*denovo* assembled and evaluated 528,259 transcripts of *C. papaya* to generate 98358 primary CDSs (Table S10). BUSCO analysis of these assembled primary CDSs (using transcriptome mode) revealed a completeness score of 86.8% which is 6% higher than the completeness score of the *C. papaya* representative genome (80.8%). However, the assembled transcriptome showed a higher duplication rate of BUSCOs (31.8%) than the reference genome (2.6%, Figure 6A), which is typical of transcriptome assemblies.

To further elucidate the differences between CDS annotations from the *C. papaya* representative genome (26,630 CDS) and our assembled 98,358 CDSs, we used a plant-specific functional annotation pipeline, Mercator4 (v5.0; Lohse et al., 2014; Schwacke et al., 2019) to annotate sequences in both sets of CDSs (Table S11 and S12; see supplementary methods S7). Out of a total of 5783 functional categories (i.e. MapMan bins; (Schwacke et al., 2019, p. 4), CDSs belonging to 5,291 bins can be found in both sets of CDSs (Common bins; Figure 6B; Table S13), indicating that the reference and assembled CDSs are largely in agreement. However, CDSs belonging to 124 bins were found exclusively in the representative genome annotation, while 267 bins were exclusive to assembled CDSs. This implies that while *de novo* transcriptome assembly can recover more protein-coding transcripts than a low-quality /partial genome, the assembly might miss some protein-coding transcripts that are lowly expressed, conditionally expressed, or silent.

To showcase the utility of transcriptome atlases generated by LSTrAP-denovo, we used the expression matrix generated by the GetExpressionMatrix.py module for co-expression analysis. We found two CDSs (*c12_49301.p1 and c0_34748.p1*) coding for phytoene desaturases (PDS; MapMan bincode: 9.1.6.1.2) present in our assembled CDSs that are missing in CDS annotations. The phylogenetic analysis featuring 11 PDSs from 9 species (Table S17; see supplementary methods S7) of different evolutionary lineages across *Viridiplantae* (the green lineage), revealed that the two PDSs share the largest evolutionary similarity with PDSs from Eudicots*, A. thaliana* and *Glycine max* (*G. max*; soybean).

Using one of the PDS *(c12_49301.p1)* as a “bait-gene” (Usadel et al., 2009) we identified the top 150 CDSs most co-expressed with *c12_49301.p1* (Table S15). This co-expression neighborhood is strongly enriched for genes involved in secondary metabolism (BH-corrected p-value < 0.0001; Table S16; Figure 6D). Further dissection of this neighborhood revealed that *c12_49301.p1* is co-expressed with members of the carotenoid biosynthetic pathway (Figure 6D) and (methylerythritol phosphate) MEP pathway. These pathways are known to be co-regulated transcriptionally given that the MEP pathway generates the precursors needed for carotenoid biosynthesis (Stanley & Yuan, 2019). Thus, it is likely that the other CDSs present in the neighborhood have a function related to carotenoid biosynthesis (Delli-Ponti et al., 2020; Lim, Zheng, et al., 2022; van Dam et al., 2018). For instance, the enzymes within the neighborhood may be accessory enzymes for the derivatization of carotenoids in *C. papaya* (Figure 6D). As such, through a simple co-expression workflow, we are able to exemplify the potential of expression data generated by LSTrAP-*denovo* in finding functionally related CDSs, and by extension, facilitating prediction/hypothesis generation of gene function in species without any genomic references.

## 5. Conclusion

The concept of using public RNA-seq data in *de novo* transcriptome assembly is not new and there exist computational pipelines developed specifically to use RNA-seq accessions downloaded from INSDC databases for de novo transcriptome assembly (Gilbert, 2019; Ortiz et al., 2021). For instance, the SRA2genes pipeline (http://arthropods.eugenes.org/EvidentialGene/other/sra2genes_testdrive/sra2genes4v_testdrive) enables users to initiate *de novo* transcriptome assembly of public RNA-seq data by specifying SRA identifiers. However, the challenge of selecting the appropriate RNA-seq accessions for the dataset remains an issue in practice, as previously alluded to in the introduction. While it is possible to devise an approach to select for RNA-seq accessions from multiple experimental conditions and sample types via annotations inferred from uploaded metadata descriptions (as seen in Goh & Mutwil, 2021), these annotations might not be accurate and available for all accessions due to the lack of format standardization and fixed vocabulary imposed on uploaded metadata. In addition, experimental conditions and sample types applicable to describe RNA-seq accessions can differ significantly from species to species, making the development of said approach difficult. As such, we have developed a novel strategy to facilitate the unbiased and automated selection of RNA-seq accessions for *de novo* assembly, and incorporated this strategy into an easy-to-use pipeline (LSTrAP-*denovo*) to generate comprehensive transcriptome atlases for eukaryotic species without genomes. As publicly accessible transcriptomic data continue to accumulate for these species, we envision that methods to interrogate and analyze said data like LSTrAP-*denovo*, will become increasingly popular, spurring the development of more advanced bioinformatics tools (e.g. *de novo* transcriptome assemblers, alignment algorithms). LSTrAP-*denovo* aims to be an open-source platform for continued development as is reflected in its extensive documentation (https://github.com/pengkenlim/LSTrAP-denovo/wiki/1.-Overview) and modular structure which can be easily upgraded by the bioinformatics community.

