## Supplemental methods and figures for "LSTrAP-*denovo*: Automated Generation of Transcriptome Atlases for Eukaryotic Species Without Genomes"

### Supplemental Figures

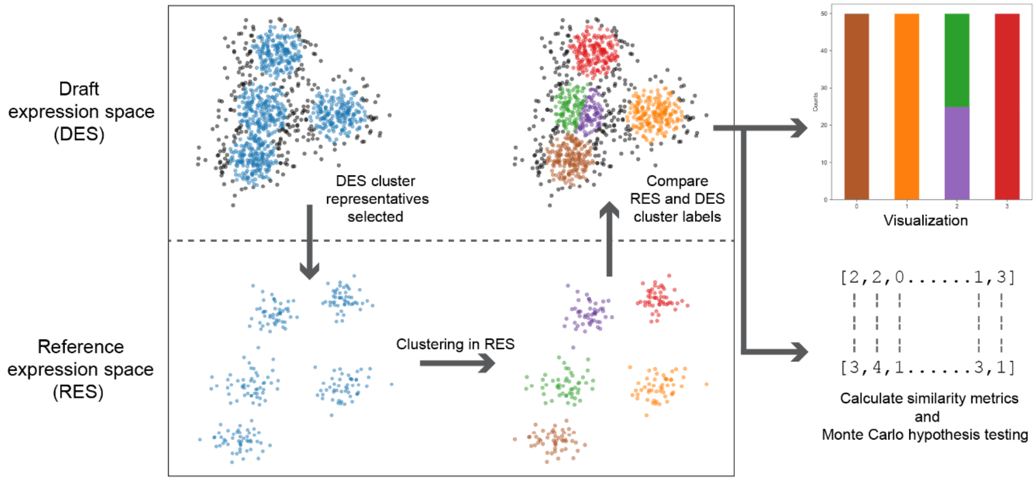


**Figure S1. K-medoids clustering method to compare DES and RES embeddings of representative RNA-seq accessions.** RES and DES are represented as 2D spaces (for simple visualization) within the box and are separated by a gray dotted line while scattered data points correspond to individual RNA-seq accessions. Accessions colored blue are representative RNA-seq accessions of clusters (25% of accessions closest to medoids), while the rest of the accessions are colored black. Brown, orange, purple, green, and red points, and bars correspond to different cluster labels assigned to representative RNA-seq accessions after K-medoids clustering.

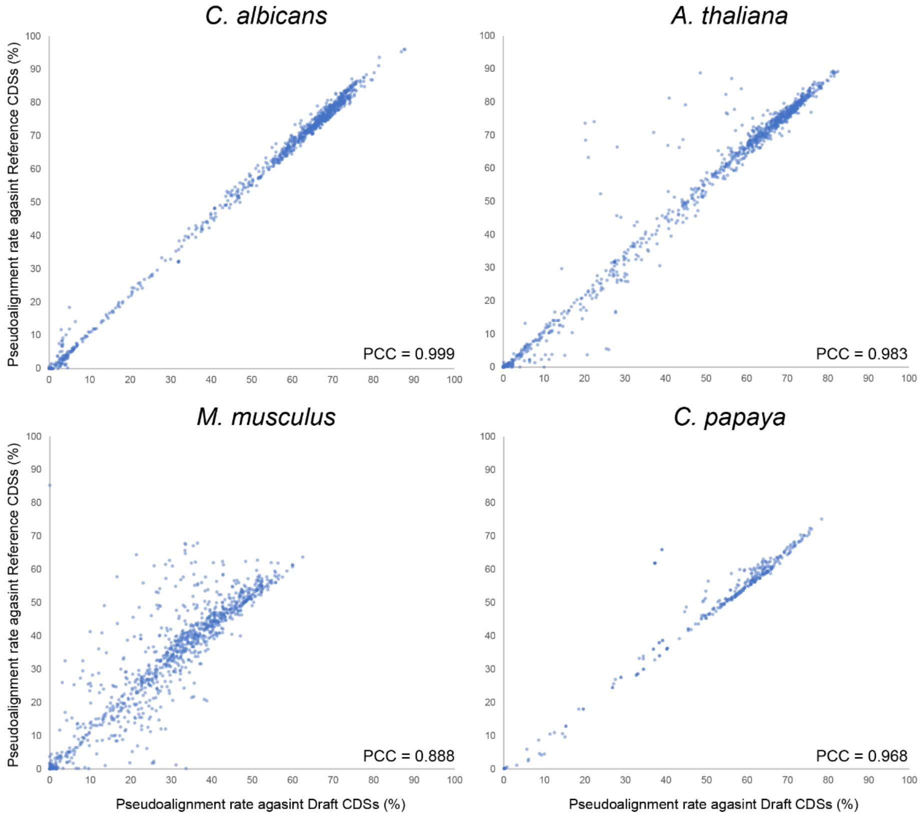


**Figure S2.Correlation of accession pseudoalignment rates when pseudoaligned against Draft CDSs and the References CDSs.** Pearson correlation coefficient (PCC) of data points calculated using linear regression are represented within each plot.

**Supplemental Methods (Pipeline Evaluation)**

**S1. Running LSTrAP-*denovo* on several test species**
The first two modules of LSTrAP-*denovo* (i.e. MakeDraftCDS.py and SelectAccessions.py) were used to process RNA-seq accessions for model species with representative genomes available on NCBI: *Arabidopsis thaliana* (*A. thaliana;* TaxID: 3702), *Mus musculus* (*M. musculus;* TaxID:10090) and *Candida albicans* (*C. albicans;* TaxID: 5476) (Table S17). Due to the large number of RNA-seq accessions for these species that are available on ENA (for e.g., >70,000 for *A. thaliana*), only a subset of RNA-seq accessions (approximately 1000) was analyzed using the `--accessions_limit 1000` flag when running SelectAccessions.py (Table S18). In addition, we also analyzed a species with poor genome quality, *Carica papaya* (*C. papaya;* TaxID: 3649) (Table S18).

**S2. Evaluation of assembled transcripts and establishment of reference CDSs**
We used the "DE novo TranscriptOme rNa-seq Assembly with or without the Truth Evaluation" (DETONATE v1.11; Li et al., 2014) workflow for the reference-based evaluation of CDSs / ORFs extracted from assembled transcripts. First, the BLAT (Kent, 2002) alignment tool was used to align CDSs / ORFs against reference CDSs and vice versa, as outlined in (https://deweylab.biostat.wisc.edu/detonate/ref-eval.html). The alignment files were then fed into the REF-EVAL module from DETONATE to calculate precision, recall, and F1 scores at the contig and nucleotide levels (Li et al., 2014). For the calculation of contig level scores, the default minimum fraction identity of 0.99 (--min-frac-identity 0.99; see https://deweylab.biostat.wisc.edu/detonate/ref-eval.html) was used. CDS annotations from representative genomes designated by NCBI genomes were used as reference CDSs; The representative genomes used were TAIR10.1 (GCF_000001735.4) for *A. thaliana*, GRCm39 (GCF_000001635.27) for *M. musculus*, Papaya1.0 (GCF_000150535.2) for *Carica papaya* (*C. papaya*) and ASM18296v3 (GCF_000182965.3) for *Candida albicans* (*C. albicans*). BUSCO v5.4 (Manni et al., 2021) analysis (transcriptome mode) was used to evaluate the completeness of *C. papaya* primary CDSs (cds_from_primary_transcripts.fasta) generated by LSTrAP-denovo using the *Brassicales* (odb10) BUSCO dataset.

**S3. Fetching of RNA-seq accessions metadata and annotation**
Title metadata of RNA-seq accessions were fetched using the *ffq* tool (Gálvez-Merchán et al., 2023). RNA-seq accessions were then annotated with tags that described their possible unsuitability for transcriptome assembly (e.g., contamination of exogenous material or sequencing library of non-coding RNA; Table S7) by searching for certain keywords within their title metadata.

**S4. Selection of representative RNA-seq accessions and generation of RES embeddings**
For each species, a subset of analyzed RNA-seq accessions was chosen as representative RNA-seq accessions. They comprise the top 25% of RNA-seq accessions that are closest to their respective cluster medoids in the k-medoids clustering carried out by LSTrAP-*denovo* as described in 2.3 (Figure S1).

To generate embeddings of representative RNA-seq accessions in reference expression space (RES), representative accessions were first pseudoaligned against reference CDSs respective to their species (see 3.3) to generate species-specific TPM expression matrices (similar to 2.2). Next, the expression matrices were normalized, and transformed into feature matrices using PCA (as described in 2.3) to generate RES embeddings of representative accessions.

**S5. K-medoids clustering of cluster representatives in RES**

Representative RNA-seq accessions is subjected to K-medoids clustering (Mohammed & Abdulazeez, 2017; Van der Laan et al., 2003) based on RES embeddings (RES clustering; Figure S1). In order to keep RES clustering of these accessions independent from DES clustering, the expected number of clusters (k) to adopt was determined based on applying K-medoids clustering through a k range of 5 to 20 (same as the default k range in *LSTrAP-denovo* used for DES clustering) and selecting the clustering iteration with the highest silhouette coefficient (Rousseeuw, 1987) instead of using the k that was previously determined for DES clustering.

**S6. Hypothesis testing and comparison of DES and RES clusterings of representative RNA-seq accessions**

To evaluate the similarity between DES and RES clusterings, we calculated the homogeneity (H) and completeness (C), as well as V-measure (V) which is derived from the harmonic mean of H and C score (Rosenberg & Hirschberg, 2007).

To evaluate the statistical significance of the observed similarity between both clusterings, we used a Monte Carlo-based method for hypothesis testing. First, outputs from both clusterings are each represented as a vector of cluster labels of size *n* (Figure S1), where the cluster label at each position corresponds to a particular observation (representative RNA-seq accession) and *n* is the number of representative RNA-seq accessions. Using the two vectors, a V score (original V score) is calculated (ref). Next, permutations of the RES clustering vector are generated via the random shuffling of cluster labels. For each permutation, a new V score is calculated based on the permuted RES clustering vector and the unchanged DES clustering vector. This facilitates the generation of a p-value for the null hypothesis where the observed similarity between the two clusterings is random. The p-value is defined as the proportion of permutations that yielded a V score that is equal to or greater than the original V score. The equation of p-value is as follows:

$$p - value = \frac{\sum_{n=1}^{N} I(V_{permuted} \geq V_{orignal})}{N}$$

where *N* is the number of permutations, *I* is an indicator function that takes a value of 0 or 1 if the statement is false or true, respectively.

**S7. Annotation of *C. papaya* CDSs and phylogenetic analysis of assembled *C. papaya* phytoene desaturases**

The primary CDSs of *C. papaya* assembled by LSTrAP-*denovo* and *C. papaya* reference CDSs were assigned Mapman functional annotations (Table S11 and S12) (Schwacke et al., 2019) using Mercator4 v5.0 ([https://www.plabipd.de/portal/web/guest/mercator4](https://www.plabipd.de/portal/web/guest/mercator4)). CDSs coding for phytoene desaturases was identified using the "9.1.6.1.2" Mapman bincode with the description of "Secondary metabolism.terpenoids.carotenoid biosynthesis.carotenes.phytoene desaturase *(PDS)".

11 phytoene desaturase protein sequences from 9 species across *Viridiplantae* used in the phylogenetic analysis were obtained from the NCBI protein database (Table S14). They are NP_193157.1 from *A. thaliana*, NP_001236769.2 from *Glycine max* (*G. max*), XP_010229080.1 from *Brachypodium distachyon* (*B. distachyon*), PTQ31715.1 from *Marchantia polymorpha* (*M. polymorpha*), GAQ82188.1 from *Klebsormidium nitens* (*K. nitens*), XP_002988989.1 from *Selaginella moellendorffii* (*S. moellendorffii*), PNW74792.1 from *Chlamydomonas reinhardtii* (*C. reinhardtii*), XP_002184512.1 from Phaeodactylum tricornutum (*P. tricornutum*), XP_002180171.1 from *P. tricornutum*, XP_003055146.1 from *Micromonas pusilla* (*M. pusilla*) and XP_003063922.1 from *M. pusilla*.

The phylogenetic tree was constructed by the neighbor-joining (NJ) method (Saitou & Nei, 1987) using the MEGA11 program (Tamura et al., 2021) based on multiple sequence alignment using the Clustal W algorithm (Thompson et al., 1994). Bootstrapping (1000 replicates) was implemented to obtain branch confidences, evolutionary distances were computed using the *p*-distance method (Nei & Kumar, 2000) and positions containing gaps and missing data were eliminated using the complete deletion option.

**S8. Co-expression analysis of C. papaya**

Columns corresponding to RNA-seq accessions with a poor pseudoalignment rate (< 20%) were removed from the TPM expression matrix generated by GetExpressionMatrix.py (Table S19; see 2.6). To calculate co-expression between a pair of CDS, Pearson's correlation coefficient (PCC) was calculated from TPM values across different RNA-seq accessions within the expression matrix subset (Usadel et al., 2009).

Network visualization of the co-expression neighborhood of *c12_49301.p1* (Table S15) was generated using Cytoscape V3.8.2 with the circular layout. Statistical tests to identify overrepresented MapMan functional categories (i.e. MapMan first-level bins) of the co-expression neighborhood were conducted via the hypergeometric test (Kemp & Kemp, 1956); null-hypothesis p-values determined for all categories were corrected for multiple-testing using the Benjamini-Hochberg method (Benjamini & Hochberg, 1995).
